# Engineering of new-to-nature halogenated indigo precursors in plants

**DOI:** 10.1101/262428

**Authors:** Sabine Fräbel, Bastian Wagner, Markus Krischke, Volker Schmidts, Christina M. Thiele, Agata Staniek, Heribert Warzecha

## Abstract

Plants are versatile chemists producing a tremendous variety of specialized compounds. Here, we describe the engineering of entirely novel metabolic pathways *in planta* enabling generation of halogenated indigo precursors as non-natural plant products.

Indican (indolyl-β-D-glucopyranoside) is a secondary metabolite characteristic of a number of dyers plants. Its deglucosylation and subsequent oxidative dimerization leads to the blue dye, indigo. Halogenated indican derivatives are commonly used as detection reagents in histochemical and molecular biology applications; their production, however, relies largely on chemical synthesis. To attain the *de novo* biosynthesis in a plant-based system devoid of indican, we employed a sequence of enzymes from diverse sources, including three microbial tryptophan halogenases substituting the amino acid at either C5, C6, or C7 of the indole moiety. Subsequent processing of the halotryptophan by bacterial tryptophanase TnaA in concert with a mutant of the human cytochrome P450 monooxygenase 2A6 and glycosylation of the resulting indoxyl derivatives by an endogenous tobacco glucosyltransferase yielded corresponding haloindican variants in transiently transformed *Nicotiana benthamiana* plants. Accumulation levels were highest when the 5-halogenase PyrH was utilized, reaching 0.93 ±0.089 mg/g dry weight of 5-chloroindican. The identity of the latter was unambiguously confirmed by NMR analysis. Moreover, our combinatorial approach, facilitated by the modular assembly capabilities of the GoldenBraid cloning system and inspired by the unique compartmentation of plant cells, afforded testing a number of alternative subcellular localizations for pathway design. In consequence, chloroplasts were validated as functional biosynthetic venues for haloindican, with the requisite reducing augmentation of the halogenases as well as the cytochrome P450 monooxygenase fulfilled by catalytic systems native to the organelle.

Thus, our study puts forward a viable alternative production platform for halogenated fine chemicals, eschewing reliance on fossil fuel resources and toxic chemicals. We further contend that *in planta* generation of halogenated indigoid precursors previously unknown to nature offers an extended view on and, indeed, pushes forward the established frontiers of biosynthetic capacity of plants.

**Graphical abstract**

**Highlights:**

- Entirely novel combinatorial pathway yielded new-to-nature halogenated indigoids.
- An array of specific chloro- and bromoindican regioisomers was generated.
- Significant retrieval rates afforded unambiguous metabolite identification.
- Engineered plants emerge as alternative manufacture chassis for fine chemicals.

## 1. Introduction

The ability of plants to produce an enormous variety of complex chemical structures is founded upon a substantive set of branched and matrix biosynthetic pathways (Fischbach and Clardy, 2007) and rooted in an almost indefinite supply of photosynthetic energy. Recent years have seen tremendous success not only in elucidation of several plant metabolic trails (Staniek *et al*., 2014) but also in their re-establishment in heterologous hosts. However, current metabolic engineering efforts are, to a great extent, aimed at grafting the pathways to fermentable recipient organisms, like bacteria or yeast (Staniek *et al*., 2014). Especially the latter, as a eukaryotic host, has been developed to accommodate plant specialized metabolite production. Conducive to activity requirements of many proteins involved in the build-up of the complex carbon skeletons of interest, the Eukaryote provides a favorable catalytic environment for, *e.g.*, the ubiquitous cytochrome P450 monooxygenases, which are anchored to the endoplasmic reticulum (ER) membrane and need to interact with their corresponding P450 reductases. Elaborate compounds of plant origin successfully produced in *Saccharomyces cerevisiae* include the sesquiterpene, artemisinic acid (Ro *et al*., 2006), the monoterpenoid indole alkaloid (MIA) precursor, strictosidine (Brown *et al*., 2015), or the benzylisoquinoline alkaloid intermediate, reticuline (Hawkins and Smolke, 2008). However, besides integration of biosynthetic genes, effective generation of plant metabolites in yeast necessitates substantive pathway engineering for precursor supply and, in most cases, retrieval rates are low, especially when compared to the plant-produced ingredients.

In contrast, recent work has shown that plants, while harboring orthologous metabolic routes, can serve as suitable vehicles for the expression of complex biosynthetic pathways (Staniek *et al*., 2013). Transient expression in host plants like *Nicotiana benthamiana* enables not only time-efficient assessment of pathway iterations but also production of novel compounds in quantity (gram-scale), as demonstrated for a suite of triterpenoids (Reed *et al*., 2017). To further investigate the potential of plant metabolic engineering, we ventured to assemble a largely artificial biosynthetic route yielding new-to-nature compounds with various applications. To attain our objective, we relied on the combinatorial use of enzymes from different sources and drew on the unique compartmentation of plant cells.

Indigoid precursors are common reagents applied in biochemical and tissue culture detection protocols. They consist of an indoxyl (3-hydroxyindole) aglycon coupled to a variable sugar moiety. Mostly colorless and soluble in aqueous solvents, when subjected to enzymatic cleavage, the reagents release reactive indoxyl, which dimerizes under atmospheric oxygen into an insoluble, deeply colored indigo dye. Aside from natural indigo precursors present in plants (Gilbert and Cooke, 2001) and mollusks (Benkendorff, 2013), several artificial halogenated derivatives were developed to vary and enhance coloration. The most prominent example is 5-bromo-4-chloro-3-indolyl-β-D-galactopyranoside, commonly known as X-Gal, used extensively in molecular biology for the blue-white screening of recombinant bacteria expressing a β-galactosidase gene (Kiernan, 2007). Currently, indigoid precursors are synthesized chemically. The need for regiospecific halogenation of aromatic carbon molecules and the low reactivity of the hydroxyl function of indoxyl make the process rather difficult (Böttcher and Thiem, 2015). Therefore, a biosynthetic alternative would be beneficial, additionally limiting reliance on organic solvents and hazardous chemicals.

Flavin-dependent halogenases, as catalysts of regiospecific halogen integration into a wide range of substrates, have been studied extensively (van Pee and Patallo, 2006; Kling *et al*., 2005). Their functionality was proposed to be based on electrophilic substitution for arene-halogenation, utilizing halide ions and oxygen (Dong *et al*., 2005; Payne *et al*., 2015). Particularly tryptophan halogenases have already found application in expanding the natural diversity of specialized compounds through integration of halogen moieties at specified positions within their structural scaffolds, greatly enhancing either pharmacological properties of a given molecule or enabling selective functionalization in subsequent chemical derivatizations. For example, introduction of tryptophan 7-halogenase PrnA into *Streptomyces coeruleorubidus* resulted in generation of a chlorinated derivative of the antibiotic pacidamycin, paving the way for its subsequent chemical modification (Deb Roy *et al*., 2010). Further, naturally halogenated antimicrobials, like rebeccamycin, were successfully diversified *via* incorporation of halogenases of divergent regioselectivity (*e.g.*, PyrH and ThaL) into the natural producer to obtain novel compounds by virtue of combinatorial biosynthesis (Sanchez *et al*., 2005). Finally, halogenases of mainly prokaryotic origin were introduced into eukaryotic systems to enable formation of entirely new compounds; upon integration of bacterial RebH and PyrH in the plant host *Catharanthus roseus*, halogenated derivatives of the endogenous monoterpenoid indole alkaloids were detected (Runguphan *et al*., 2010; Glenn *et al*., 2010). The highlighted work shows that tryptophan halogenases are versatile tools for engineering of novel biosynthetic routes, also in plant metabolism.

In this study, we report successful production of a set of halogenated indigoid precursors in a plant-based system, providing a biotechnological alternative to chemical synthesis from fossil fuel resources and highlighting the potential of plants as sustainable chassis for the manufacture of useful halogenated fine chemicals.

## 2. Results and discussion

### 2.1 Expanding an established indican pathway did not yield halogenated derivatives

As we were expecting to test a large number of iterations of biosynthetic routes, encompassing numerous catalyst genes with diverse signal sequences for subcellular targeting as well as variable regulatory elements, we applied the modular cloning technique GoldenBraid (Sarrion-Perdigones *et al*., 2013; Patron *et al*., 2015; Fig. 1). The relevant expression constructs, mobilized into *Agrobacterium tumefaciens,* afforded transient transformation of *N. benthamiana* leaves.

**Figure 1.**
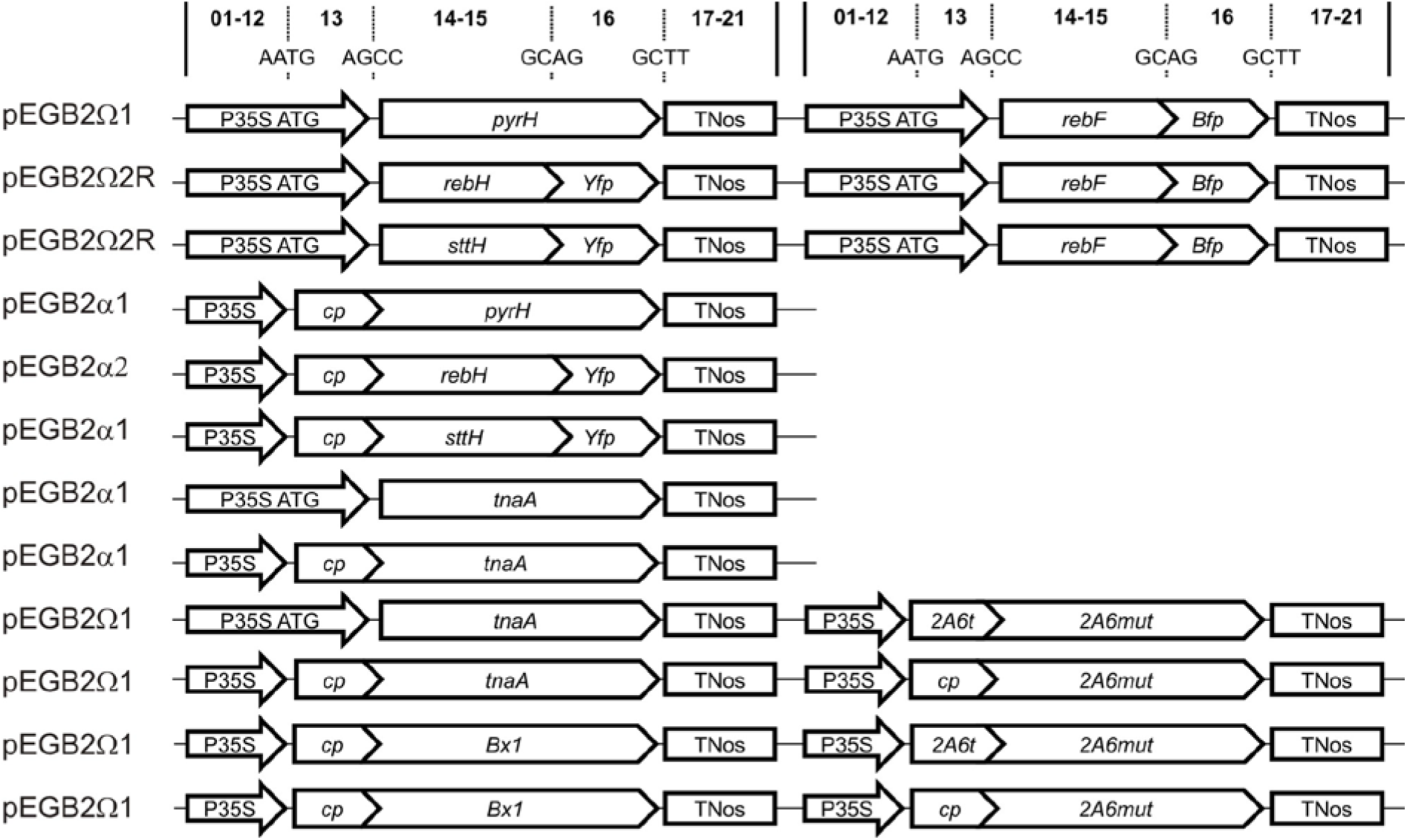
Schematic representation of the expression constructs used in the study. Numbers on top of the diagram represent standard GoldenBraid classes within the TU structure, while capital letter designations indicate four-nucleotide overhangs ascertaining seamless assembly of GBparts (Sarrion-Perdigones *et al*., 2013). P35S, cauliflower mosaic virus (CaMV) 35S promoter; P35S ATG, CaMV 35S promoter with the integrated start codon (spanning GB positions 01–13); *2A6t*, signal peptide sequence of the P450 2A6 monooxygenase; *cp*, chloroplast transit peptide sequence; TNos, terminator of the nopaline synthase gene from *A. tumefaciens*. For disambiguation of gene name abbreviations, see main text below

To evaluate the potential to generate halogenated indigoid precursors, we initially chose to build upon a previously reported, partially orthologous indican (indolyl-β-D-glucopyranoside) biosynthetic pathway (Warzecha *et al*., 2007). We showed that the combination of genes encoding for indole synthase (Bx1) from maize and the human cytochrome P450 monooxygenase 2A6 yielded the indigoid in stably transformed *Nicotiana tabacum* plants. This artificial pathway diverts from tryptophan biosynthesis in that it utilizes its direct precursor, indole 3-glycerolphosphate to form indole, which is then converted to indoxyl and, further on, to indican by endogenous tobacco glucosyltransferases (Fig. 2).

**Figure 2.**
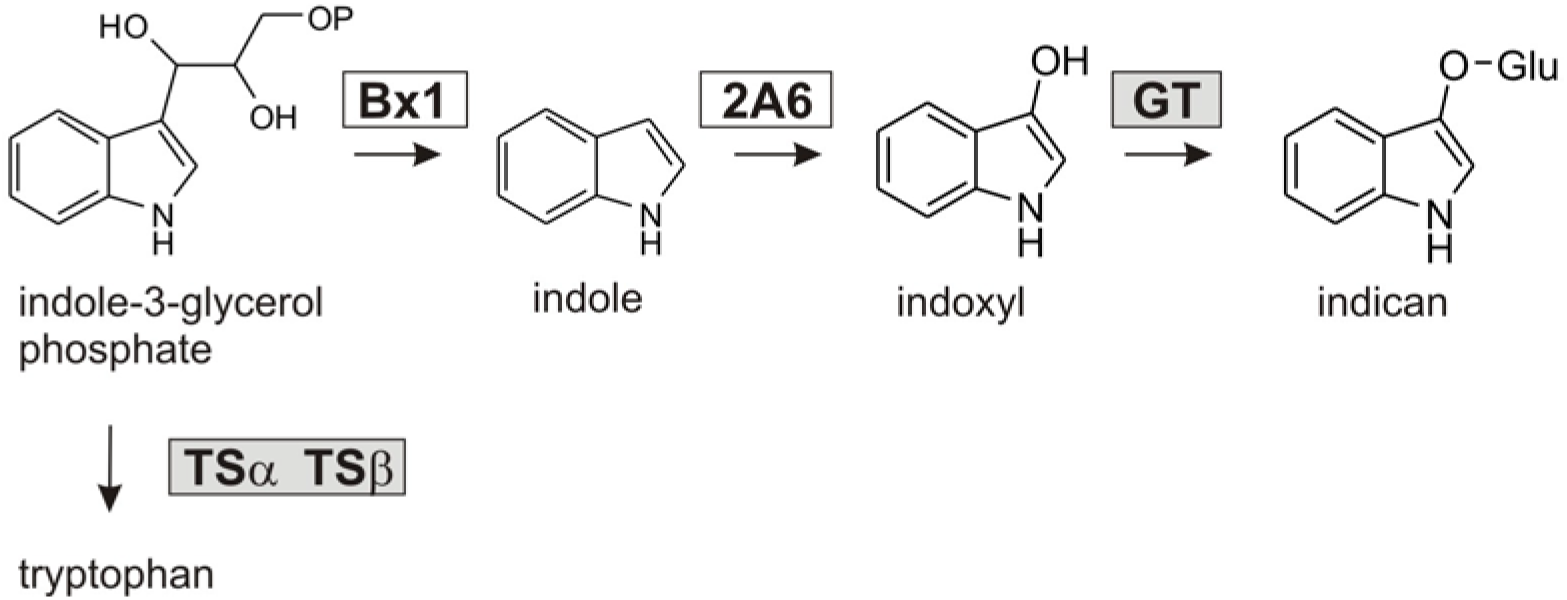
Previously reported synthetic pathway leading to indican production in tobacco plants (Warzecha *et al*., 2007). Bx1, indole synthase from *Z. mays*; 2A6, human cytochrome P450 monooxygenase 2A6; enzymes endogenous to the host plant (shaded boxes),TS, tryptophan synthase and GT, glucosyltransferase

Engineering of the proposed pathway towards halogenation would likely spur generation of substituted indole, which might have a hampering effect on its subsequent conversion by P450 2A6. To facilitate the latter step, we decided to employ not the native human cytochrome but its mutant exhibiting elevated affinity for substituted indole derivatives, amongst them 5-, 6-, and 7-halogenated variants (Nakamura *et al*., 2001). Accordingly, we integrated the P450 monooxygenase 2A6 mutant L241C/N297Q, further dubbed 2A6mut, into our pathway build-up.

Our next objective was to determine the stage in the engineered pathway most favorable for the introduction of the halogen (chlorine under physiological conditions) into the indole moiety. Here, we selected three bacterial halogenases shown to chlorinate tryptophan at various positions, namely PyrH (from *Streptomyces rugosporus*, halogenating Trp at C5), SttH (from *Streptomyces toxitricini*, generating 6-chlorotryptophan), as well as RebH (Trp 7-halogenase from *Lechevalieria aerocolonigenes*). All three halogenases were previously reported functional *in planta* (Runguphan *et al*., 2010; Fräbel *et al*., 2016). To facilitate their activity, we also employed the NADH-dependent flavin reductase RebF, responsible for providing reducing equivalents to the halogenases in bacteria and in the plant cell cytosol (Fräbel *et al*., 2016).

A possible route to haloindican in the proposed setting would require coupling of the halogen substrate to indole or indican but not tryptophan, the latter not being directly involved in the biosynthetic trail. To exclude the halogenation of indican itself, we infiltrated leaves expressing the halogenase (PyrH, SttH, and RebH) and reductase (RebF) genes with its 1 mM solution. However, the supplementation did not spur chloroindican formation in any of the three transient systems, while indican concentrations in leaves harboring active halogenases ranged from 84.4 ± 20.3 to 63.5 ± 7.0 nmol/g FW and were comparable to those characteristic of control samples (86.0 ± 5.0 nmol/g FW; wild type leaves infiltrated with indican). Thus, the metabolite was discounted as a potential substrate of the investigated halogen-integrating catalysts. Concurrently, and in accord with our previous observations (Fräbel *et al*., 2016), considerable amounts of chlorotryptophan accumulated in the analyzed leaf tissue.

The purported substitution of indole was then examined *via* incorporation of Bx1 in concert with the halogenases (augmented by the partner NADH-dependent flavin reductase, RebF) and the P450 2A6mut into tobacco leaves. While the precursor: indole, the product: indican, as well as halogenated tryptophan were generated proving functionality of the individual genetic constructs, co-expression of the halogenase genes within the established pathway did not result in detectable levels of chlorinated indican. The setback was presumably caused by the incompatibility of the selected catalysts, driving C5, C6, and C7 halogenation, with the substrate. Had it been processed, the halogen atom would presumably substitute indole at C3, reported as the most electronically favorable position of the pathway intermediate (Payne *et al*., 2015), consequently inhibiting its P450-driven hydroxylation. In conclusion, neither indican nor indole served as a suitable intermediate for direct *in planta* halogenation by virtue of the investigated microbial enzymes.

### 2.2 Pathway redesign and integration of bacterial tryptophanase enabled formation of chloroindican *in planta*

Since tryptophan is efficiently converted by all three halogenases (Fräbel *et al*., 2016), we designed an alternative biosynthetic route combining coding sequences of the halogen-integrating catalysts and the mutant P450 2A6 with the bacterial gene encoding tryptophanase TnaA. The enzyme was shown to readily process the fundamental substrate to indole as well as accept its halogenated derivatives, like 5-, 6-, and 7-chlorotryptophan (Lee and Phillips, 1995). The amended approach would thus ensure chlorination of the amino acid by the halogenases followed by its conversion to chloroindole. The latter, serving as a substrate for the mutated monooxygenase, would form chloroindoxyl, ultimately transformed to chloroindican by virtue of ubiquitous plant glycosyltransferases (Fig. 3).

**Figure 3.**
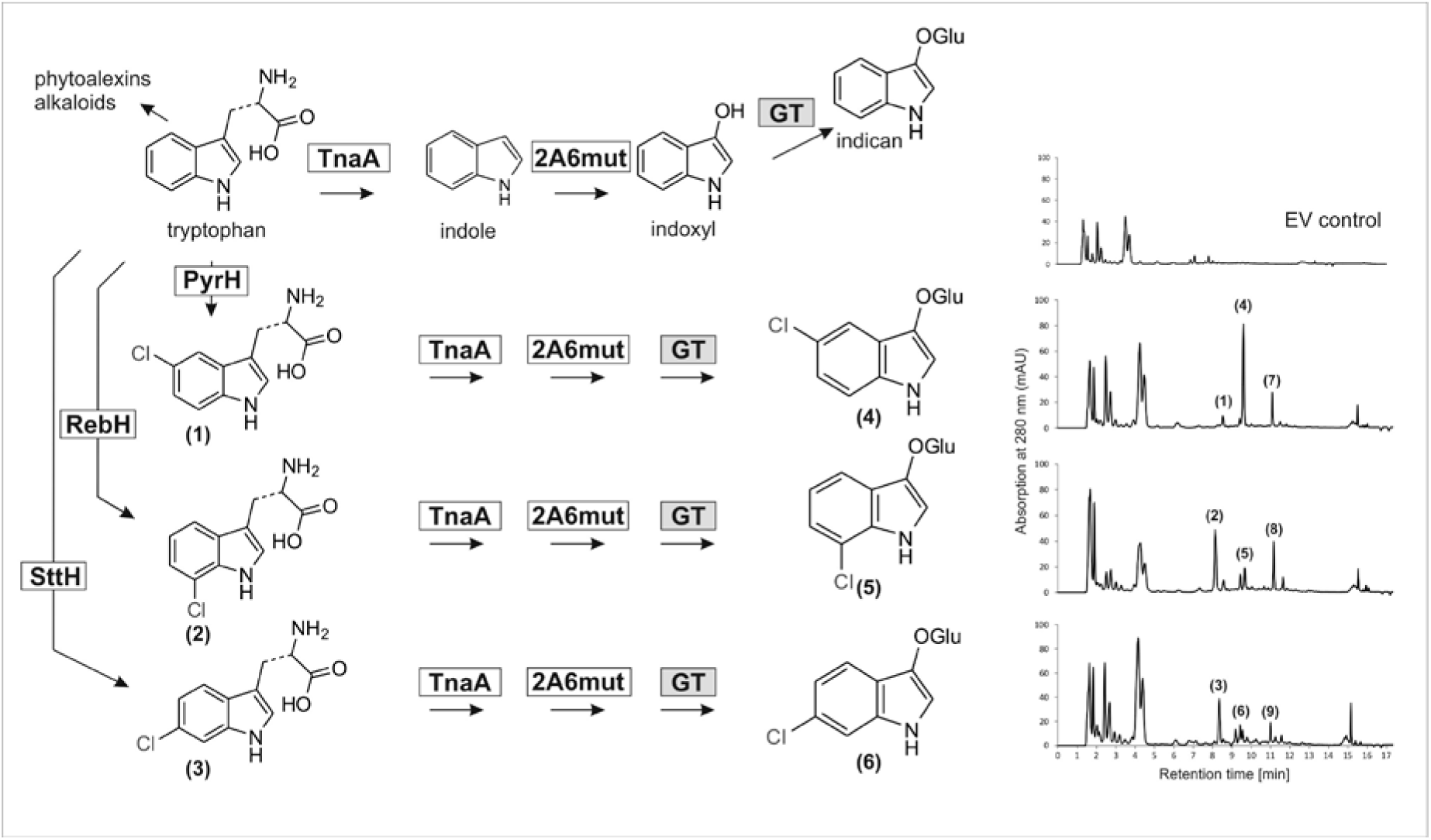
Engineered biosynthetic pathways yielding halogenated indican derivatives in tobacco. Shaded boxes, enzymes endogenous to the host plant (GT, glucosyltransferase); 2A6mut, L241C/N297Q mutant of the human cytochrome P450 monooxygenase 2A6; TnaA, tryptophanase from *E. coli*; PyrH, Trp 5-halogenase from *S. rugosporus*; SttH, Trp 6-halogenase from *S. toxitricini*; RebH, Trp 7-halogenase from *L. aerocolonigenes*; (1) 5-chlorotryptophan; (2) 7-chlorotryptophan; (3) 6-chlorotryptophan; (4) 5-chloroindican; (5) 7-chloroindican; (6) 6-chloroindican; (7) side-product of 5-chloroindican biosynthesis; (8) side-product of 7-chloroindican biosynthesis; (9) chlorinated side-product of 6-chloroindican biosynthesis; EV, empty vector

Indeed, analysis of extracts from plants infiltrated with the relevant combinatorial expression constructs corroborated formation of new metabolites (Fig. 3). Beside the chlorinated tryptophan derivatives, novel halogenated compounds, whose *m/z* values and isotope patterns were in accord with those characteristic of mono-chlorinated indican, were detected *via* liquid chromatography coupled with mass spectrometry (LC-MS; Fig. 4). Moreover, in the case of SttH, the retention time value and the mass fragmentation pattern of the newly generated product corresponded to those of the authentic 6-chloroindican standard.

**Figure 4.**
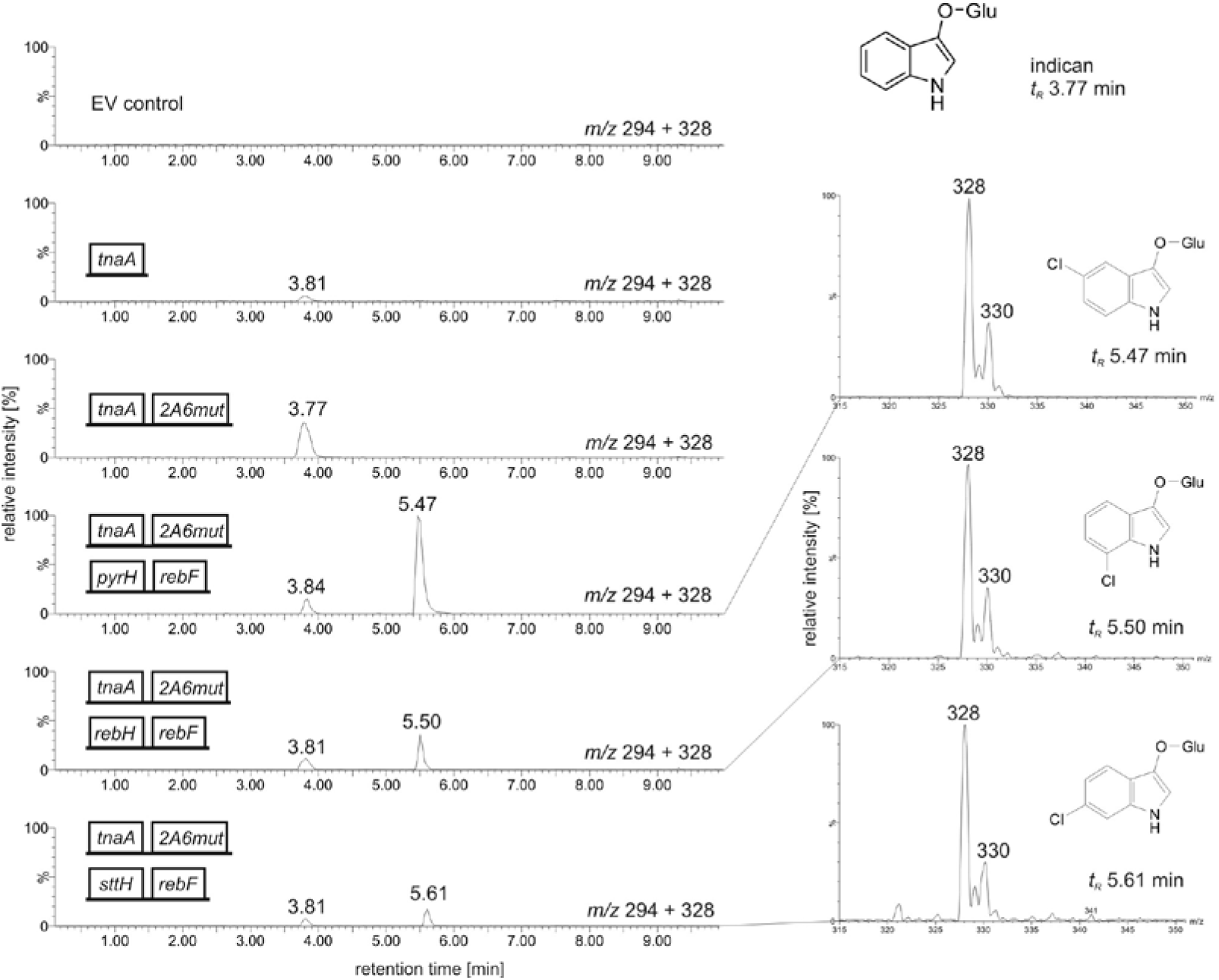
UPLC-MS analysis of engineered-plant extracts encompassing indican and chloroindican derivatives. Formation of indican was confirmed after co-localization of TnaA and 2A6mut in the cytosol. Biosynthesis of chloroindican derivatives was ascertained upon incorporation of the investigated tryptophan 5- (PyrH), 7- (RebH), and 6- (SttH) halogenases (in the cytosolic context shown here, additionally augmented by the reductase RebF) into the engineered pathway. EV, empty vector; indican: *m/z* 294, *t_R_* ~3.80 min; 5-chloroindican: *m/z* 328, *t_R_* 5.47 min; 7-chloroindican: *m/z* 328, *t_R_* 5.50 min; 6-chloroindican: *m/z* 328, *t_R_* 5.61 min. Mass analysis shows a pair of [M]^-^ ions at *m/z* 328 and 330 in a 3:1 ratio

In all samples, non-halogenated indican was detected (*t_R_* ~ 3.80 min; within range of the authentic standard, *t_R_* 3.81 min) and its presence confirmed *via* mass analysis of the relevant LC-peaks (*m/z* 294; Supplementary Fig. 1). Small amounts of the metabolite were also recorded in plants harboring TnaA alone, indicating partial conversion of indole to indoxyl by virtue of an endogenous monooxygenase. However, the heterologous host-specific activity produced but traces of indican.

With the modular build-up of the orthologous pathway, we further tested different in-cell localizations of the novel biosynthetic route. While previous studies proved that ER-targeting of the halogenases does not yield tryptophan substituents of interest (Fräbel *et al*., 2016), we channeled the relevant activities to the cytosol and chloroplasts. Both investigated intracellular contexts were validated as functional pathway expression venues, producing haloindican derivatives. Remarkably, the catalytic activity of 2A6mut was maintained upon its targeting to the plastids (Supplementary Fig. 2). The unique context of the organelle further supported halogenation of tryptophan by both SttH and RebH, rendering the co-expression of *rebF* superfluous, as previously observed for the P450 2A6 (Fräbel *et al*., 2016). The requisite reducing augmentation was either fulfilled by a plastidic ferredoxin (Fd) or acquired directly from the photosystem, as reported for various P450 monooxygenases (O’Keefe *et al*., 1994) (Lassen *et al*., 2014).

Thin layer chromatography (TLC) further confirmed the presence of indigogenic compounds in the investigated extracts (Fig. 5). While indican and its derivatives are largely colorless and not visible on silica plates under daylight, cleavage of their sugar moieties (spurred by the hydrolytic activity of the applied spraying reagent) facilitated dimerization of deglucosylated scaffolds to indigoid dyes; hence the development of characteristic blue to purple coloration.

**Figure 5.**
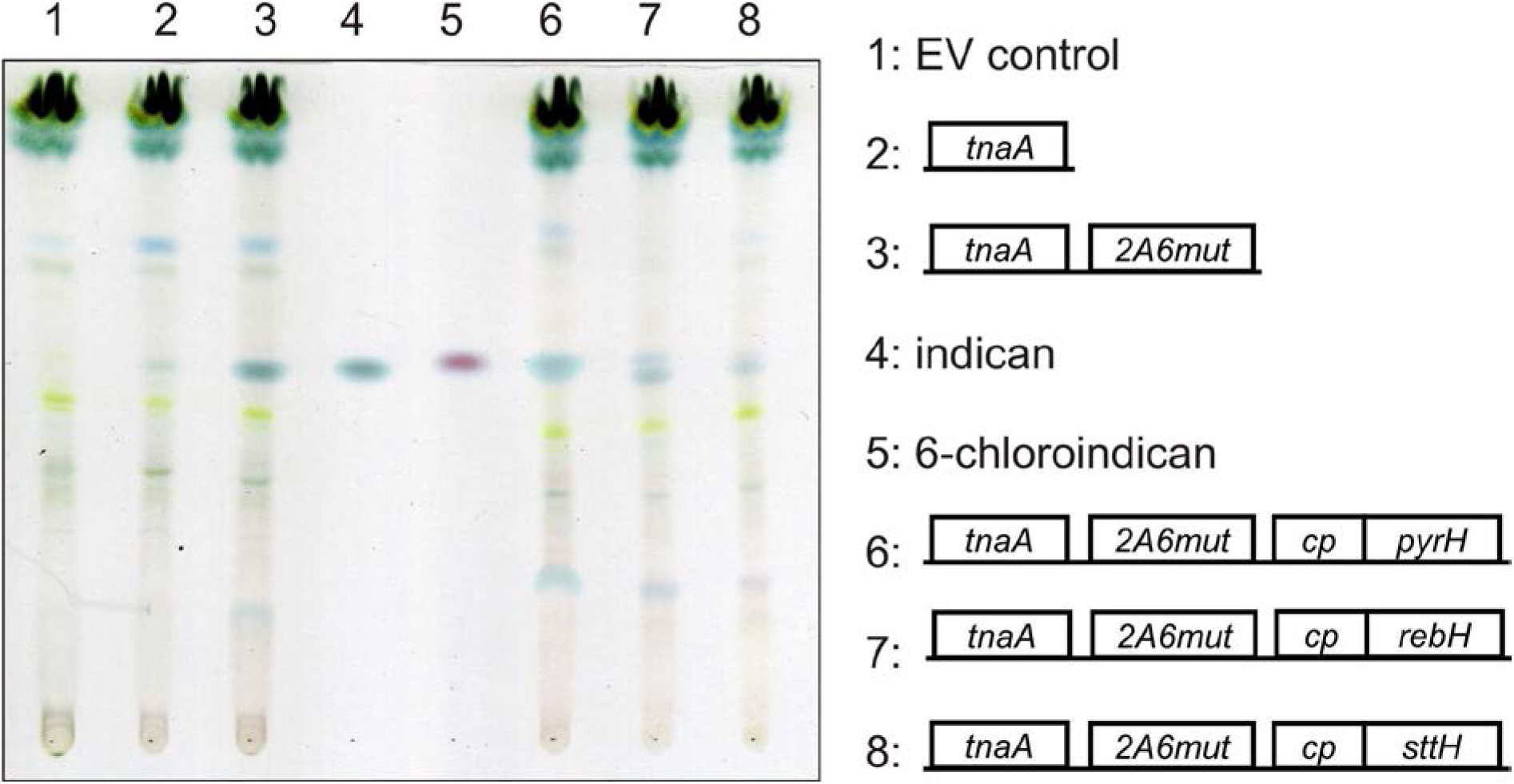
TLC analysis of extracts derived from plants engineered to produce chloroindican. Biosynthesis of chloroindican derivatives was corroborated in plants harboring TnaA and 2A6mut localized in the cytosol and either PyrH (Trp 5-halogenase), RebH (Trp 7-halogenase), or SttH (Trp 6-halogenase) localized in chloroplasts. Small amounts of indican were synthesized upon sole expression of *tnaA*, pointing to partial conversion of indole to indoxyl catalyzed by an endogenous monooxygenase. EV, empty vector; indican: *R_f_* 5.40; 5-chloroindican: *R_f_* 0.55; 7-chloroindican: *R_f_* 0.57; 6-chloroindican: *R_f_* 0.56

The TLC analysis also showed that, beside halogenated indican derivatives, the non-substituted metabolite remained present in the leaf extracts. While spraying with the hydrolyzing reagent (ethanolic HCl) efficiently uncouples the sugar moiety, the dimerization process of the released indoxyl is highly variable and, additionally, dependent on the position of the halogen substituent (Kiernan, 2007). Thus, the observed coloration intensity might not necessarily reflect the quantity of indican and its derivatives in individual samples. For this reason, metabolite quantification was performed with HPLC, using calibration curves of standard reference compounds.

### 2.3 Chloroindican accumulated in plants in significant amounts

The amounts of chloroindican obtained in the infiltration experiments involving individual halogenases differed considerably, with PyrH generating highest metabolite levels of up to 0.93 ±0.089 mg/g dry weight (DW) while RebH- and SttH-driven yields were 0.31 ±0.017 mg/g DW and 0.10 ±0.016 mg/g DW, respectively. The observed variation could stem from inconsistencies in experimental conditions inherent to the infiltration process. In a series of tests addressing the issue, introducing *Agrobacterium*-borne constructs containing halogenase genes alone, the three investigated enzymes showed similar catalytic activities *in planta* (Supplementary Fig. 3). However, when the amounts of the assorted products of the engineered pathways were compared, a different picture arose (Supplementary Fig. 3). Only in the presence of TnaA and 2A6mut acting in concert with the 5-halogenase PyrH, chlorotryptophan was efficiently converted to chloroindican (in this case, 5-chloroindican). With RebH as the halogenating catalyst, the turn-over of 7-chlorotryptophan to 7-chloroindican was less effective, as demonstrated not only by the reduced chloroindican content, as compared to the PyrH-pathway output, but also by the increased accumulation of 7-chlorotryptophan. When SttH was analyzed in the context of the entire metabolic trail, (6-)chloroindican yields were lowest. The denoted variation could, therefore, be due to disparate conversion efficiencies of the engineered pathway enzymes (TnaA and 2A6mut) against the differentially halogenated tryptophan and/or subsequent intermediates, indicating that the position of the halogen moiety within the substrate scaffolds might be the fulcrum of efficacious haloindican biosynthesis.

Our previous observations corroborated dual halogenation of tryptophan upon co-expression of *rebH* and *sttH*, resulting in retrieval of 6,7-dichlorotryptophan (Fräbel *et al*., 2016). Consequently, we tested combinations of halogenase encoding genes in our GB tool-box. However, no double- or triple-halogenated indican was detected in the experiments involving co-integration of the corresponding catalysts within the context of the entire artificial metabolic trail; only the mono-halogenated indican and tryptophan derivatives as well as di-halogenated amino acid variants were corroborated as constituents of the analyzed plant extracts. Taken together, insufficient promiscuity of the heterologous enzymes toward halogenated substrates, as compared to their parent molecules, seems to be limiting the potential diversity of products with an indican core structure. Enzyme engineering through design of mutants exhibiting broader substrate specificity might help overcome this obstacle, as has been shown, *e.g.*, for strictosidine synthase (Bernhardt *et al*., 2007; Loris *et al*., 2007). Here, single amino acid exchanges in the substrate binding pocket allowed the catalyst to accept substituted tryptamine derivatives, expanding the range of possible reaction partners and, thereby, the repertoire of formed products. One of these mutants was successfully applied *in planta* to yield halogenated monoterpenoid indole alakloids (Runguphan *et al*., 2010).

### 2.4 Identity of plant-produced 5-chloroindican was confirmed by NMR

Biosynthesis of the putative 5-chloroindican proved more efficient compared to the other regioisomers, yielding sufficient amounts for subsequent purification. Consequently, the metabolite was isolated from the relevant leaf extracts by preparative TLC and its structure was elucidated by nuclear magnetic resonance (NMR) spectroscopy. To facilitate assignment of the spectra and unambiguously determine the structure of the new compound, multidimensional correlation spectra (HSQC, HMBC, and NOESY) were recorded and compared to the reference spectra of indican and 6-chloroindican. Identification of NMR signals and their assignment (numbering scheme, Supplementary Fig. 4) was straightforward for the spectra of the reference compounds. In case of the engineered product extract however, the signals of the substituted indican were, in some cases, obfuscated due to overlap with those characteristic of impurities. Using standard stimulated echo diffusion ordered spectroscopy, we were not able to separate the resonances of the desired compound from those of the contaminants, particularly in the spectral region of the sugar moiety. Presumably, the signal overlap in the STE-DOSY resulted in multiple species with different self-diffusion constants being treated as equivalent in the monoexponential Stejskal-Tanner fit, thus showing severe distortions in the DOSY plot. The recently proposed PSYCHE-iDOSY method (Foroozandeh *et al*., 2016) aims to alleviate this shortcoming of the classic technique by broadband homonuclear decoupling of the spectrum in the ^1^H dimension. The collapse of the ^1^H multiplets to singlets reduced the signal overlap significantly and allowed for a much cleaner fit of the diffusion data (Supplementary Fig. 9), ultimately leading to the unambiguous identification of the problematic resonances as belonging to the non-naturally substituted 5-chloroindican. Subsequent assignment of its NMR data proceeded classically, through analysis of the 2D correlation spectra (Supplementary Fig. 5–8, Supplementary Table 2). The substitution position of the indole moiety is easily identifiable by virtue of the *J*-coupling pattern of the aromatic resonances and their NOE contacts to the anomeric proton (Fig. 6). Similarly, the chemical shifts and *J*-coupling pattern of the sugar moiety are easily identifiable as those of a β-glucopyranoside. Concerted application of the aforementioned methods afforded confirmation of the identity of the engineered metabolite as 5-chloro-3-indolyl-β-D-glucopyranoside (5-chloroindican). Although none of the employed techniques enables definitive determination of the absolute configuration of the glucose enantiomer present in indican derivatives, the absence of L-glucose in living organisms led us to the conclusion that the engineered compound was, indeed, the D-glucoside.

**Figure 6.**
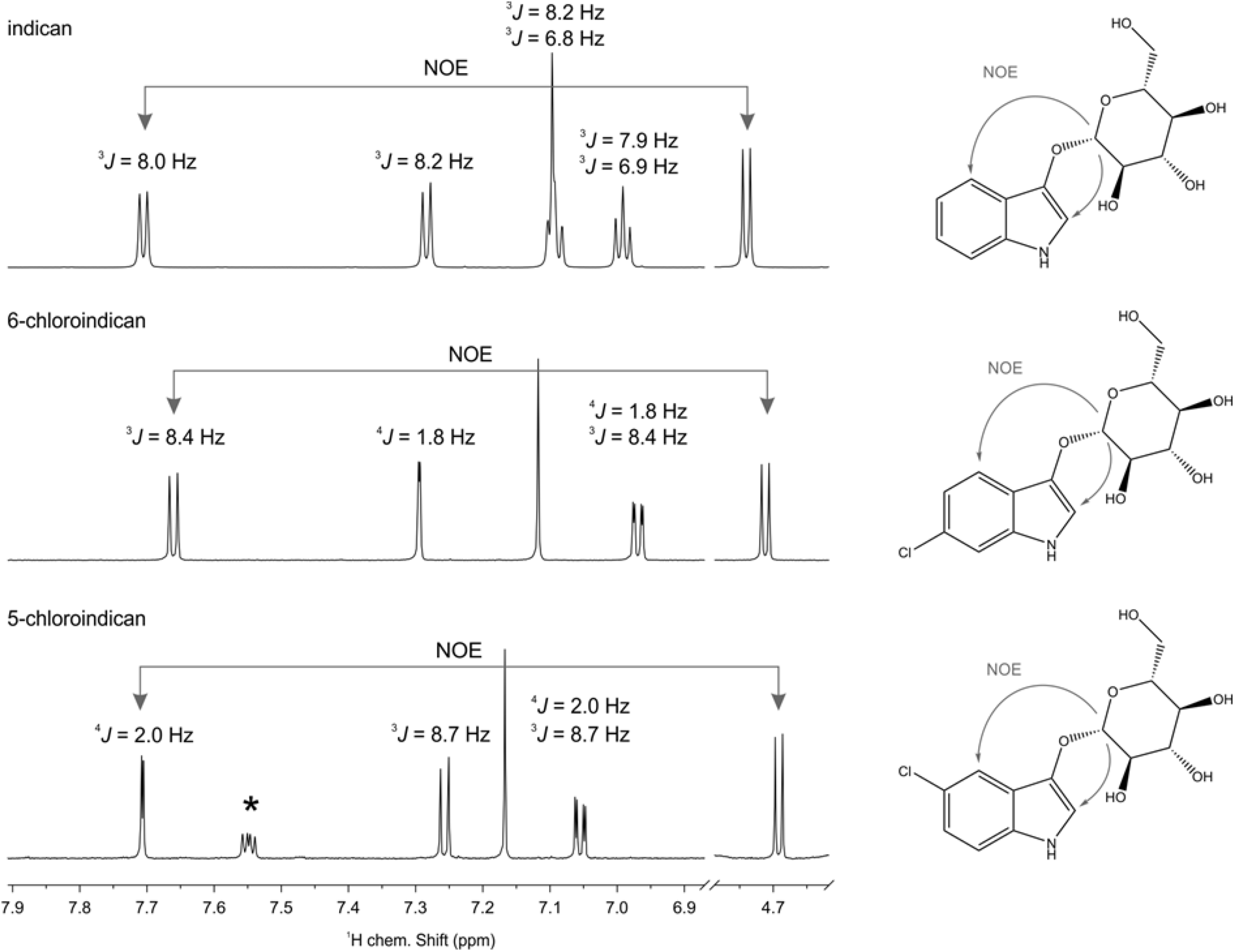
Stacked plot of the regions of interest of the ^1^H NMR spectra of indican (top panel), 6-chloroindican (middle panel) and the isolated 5-chloroindican (bottom panel). The *J*-coupling pattern of the aromatic resonances is characteristic of the substitution pattern of the indol moiety. Together with the observed NOE contacts, it enables unambiguous assignment of each regioisomer. Impurities in the plant extract are marked with an asterisk. The complete set of spectra and full assignment of all resonances are provided in the Supplementary material

### 2.5 Substrate supplementation afforded production of bromo-substituted indican

Lastly, we established the feasibility of *in planta* generation of bromo-substituted indican. While physiological conditions in plant cells favor chlorination due to relatively high chloride concentration (up to 10 mM; (Barbier-Brygoo *et al*., 2000), supplementation of bromide enables efficient bromination of tryptophan by the respective halogenases (Runguphan *et al*., 2010; Fräbel *et al.,* 2016). Therefore, *N. benthamiana* leaves were infiltrated with KBr solution (40 mM) one day after their transient transformation with the investigated pathway gene combinations. UPLC-MS analysis of the resulting plant extracts indeed corroborated formation of brominated indican derivatives corresponding to regiospecificity of each halogenase (Supplementary Fig. 10). Due to the aforementioned inherent presence of chloride in the leaves, chlorinated indican was still produced, rendering bromo-substituents only a fraction of the total indican content.

## 3. Conclusions

Our study shows that grafting artificial metabolic pathways into plants results in formation of novel, often potentially valuable metabolites. While some, like the aforementioned 5-chloroindican, might accumulate in significant amounts, additional engineering steps, *e.g.*, targeted modification of precursor supply, promise to further boost product retrieval rates. Thus, plants emerge as versatile and viable chassis for the manufacture of bespoke halogenated fine chemicals, offering an alternative to fossil fuel-reliant chemical synthesis approaches. Moreover, analysis of plant metabolite profiles upon concerted expression of the engineered pathway genes might result in detection of entirely unique compounds. As the engineering efforts perturb the endogenous plant cell biosynthetic systems, a yet untapped resource of new-to-nature specialized (side-)metabolites comes into view.

## 4. Materials and methods

### 4.1 DNA cloning

All DNA constructs used for transient transformation of *N. benthamiana* plants were assembled relying on the GoldenBraid 2.0 (GB2.0) cloning technique (Sarrion-Perdigones *et al.,* 2013) as described previously (Fräbel *et al.,* 2016). GB-customized *pyrH (S. rugosporus,* GenBank AY623051.1) was commercially synthesized (Integrated DNA Technologies, Coralville, IA, USA), while system-compatibility of the remaining genetic elements was attained by virtue of PCR. Genomic DNA of *E. coli* DH5α (one µL of cell suspension per reaction) was used for amplification of *tnaA* (Gene ID 948221), whereas a plant codon-optimized P450 2A6 L241C/N297Q mutant gene template sequence (*2A6mut*) was purchased from Mr. Gene GmbH (Regensburg, Germany). Further, *Bx1* (*Z. mays,* Gene ID 542117) (Warzecha *et al*., 2007) and *2A6mut* were integrated into the GoldenBraid cloning system without their innate targeting sequences, thus spanning positions 14–16 of the GoldenBraid TU build-up (Sarrion-Perdigones *et al*., 2013); Fig. 2). These were replaced by the standardized chloroplast transit peptide sequence (*cp*; (Fräbel *et al*., 2016) and/or a stand-alone GBpart encompassing the DNA fragment encoding for the 2A6 signal peptide (*2A6t*). The latter was generated by annealing two appropriately designed oligonucleotides (Supplementary Table 1), 40 pmol each, through incubation in 20 µL of 1× ligase buffer (Promega, Madison, USA) at 95 °C for five min followed by a two-hour cooling period at room temperature (RT). Subsequent phosphorylation reaction catalyzed by T4 polynucleotide kinase (Thermo Fisher Scientific Inc., Waltham, USA) was performed according to the manufacturer’s recommendations, and its product applied in the GoldenBraid domestication procedure. All relevant oligos are listed in Supplementary Table 1.

### 4.2 Transient gene expression in *N. benthamiana*

Transformation of plants was carried out as reported previously (Frabel et al. 2016). All infiltrations were performed in three biological replicates; in case of co-infiltration of alternative individual *A. tumefaciens* cultures, the ratios of all relevant constructs were kept constant within each experiment. For the analysis of direct indican halogenation, tobacco leaves were infiltrated with the relevant bacterial cultures (harboring RebF in tandem with PyrH, SttH, and RebH constructs, respectively). Two days post-transformation, 1 mM indican solution (in water-methanol, 19:1 v/v) or the control solution (water-methanol, 19:1 v/v) were delivered. The leaves were harvested four days after indican infiltration. To ensure reliable quantification of chloroindican derivatives and purification of 5-chloroindican, vacuum-infiltration was applied (final OD_600_ of bacterial suspensions, 0.7); plant material was harvested and processed four days after *Agrobacterium*-mediated gene transfer.

### 4.3 Chromatographic analysis of halogenated metabolites

In the main, extraction and analyses of halogenated metabolites by HPLC and UPLC-MS were executed according to the previously established procedure (Fräbel *et al*., 2016). The modifications applied herein pertained to 1) indican detection by HPLC: separation of leaf extract constituents on the Zorbax Bonus RP-C14 150×4.6 mm, 5 µm analytical column (Agilent, Santa Clara, Ca, USA) with the solvent system comprising 0.1% (v/v) formic acid (A) and methanol (B) applied in the nonlinear gradient (% B) of 0–3 min, 30; 3–5 min, 30 to 70; 5–10 min 70, and 2) mass spectrometry: detection of metabolites in the negative ESI mode with a full scan MS experiment (*m/z* 50–1,000) and a capillary voltage of three kV. Bromoindican derivatives synthesized *in planta* were analyzed by LC-MS in two biological replicates. For TLC, 200 mg of freshly ground leaf powder were extracted with one mL of 0.5% acetic acid in acetone (v/v) for 15 min at RT and centrifuged for ten min at 17,000 ×*g*. 500 µL of the supernatant were concentrated at 45 °C under vacuum and the residue was dissolved in 50 µL of methanol. Ten µL of each leaf extract were applied onto the TLC plates (ALUGRAM Xtra SIL G UV_254_, Macherey-Nagel, Düren, Germany) and their constituents separated in ethyl acetate-methanol-water-ammonia (75:15:10:0.07, v/v/v/v). The solvent composition was optimized to ensure efficient visualization of small amounts of 7- and 6-chloroindican (through controlled migration of metabolites along narrow paths) and sufficient separation of indican and chloroindican derivatives. 6-Chloro-3-indolyl-β-D-glucopyranoside (Glycosynth Limited, Warrington, UK) and 3-indoxyl-β-D-glucopyranoside (AppliChem, Darmstadt, Germany) served as standard compounds (1 mg/mL of methanol; final applied amount, 2.5 µg). Indican and its halogenated counterparts were detected upon spraying the plates with 5% HCl in ethanol (v/v) and their subsequent heating.

### 4.4 Quantification of chloroindican derivatives

To determine biosynthetic yields of each of the engineered metabolites of interest, leaves of three relevant *N. benthamiana* transformants were combined, ground in liquid nitrogen, and freeze-dried (with five to seven pooled samples analyzed subsequently). One g of freeze-dried leaf material was extracted with 20 mL of 10% methanol (v/v) and homogenized for 30 min by sonication. The crude extract was purified form solid particles by filtration using Miracloth (Merck, Darmstadt, Germany) and *via* centrifugation at 17,000 ×*g* for 10 min. Concentration values of chlorinated indican derivatives in leaf tissue were determined based on integration of HPLC peak areas characteristic of an array of serial dilutions of the 6-chloro-3-indolyl-β-D-glucopyranoside standard.

### 4.5 Purification of 5-chloroindican by preparative TLC

The relevant methanol extracts (prepared as described above) were freeze-dried, dissolved in one-tenth of the initial volume of the solvent, and centrifuged at 17,000 ×*g* for 5 min. The resultant supernatants were applied onto ALUGRAM TLC plates and the metabolites separated three times (on one plate) in ethyl acetate-methanol-water-ammonia (80:15:5:0.07, v/v/v/v). The modified solvent composition afforded optimal separation of indican and chloroindican derivatives, effectively preventing cross-contamination. A narrow (vertical) portion of each TLC plate was sprayed with ethanolic 5% (v/v) HCl and, upon halometabolite visualization, used as a reference for the identification of 5-chloroindican on the untreated plate area under UV light. The corresponding silica layer was removed (scraped with a spatula) and 5-chloroindican was eluted from the solid phase with one mL of methanol. All isolate extracts were pooled and the solvent disposed of (45 °C vacuum and air flow evaporation). The residual water was eliminated by freeze-drying.

### 4.6 Structure elucidation by NMR spectroscopy

Reference samples were prepared by dissolving five mg of the available standard compounds, 6-chloro-3-indolyl-β-D-glucopyranoside and 3-indoxyl-β-D-glucopyranoside, in 150 µL of 99.95% methanol-*d*4 (Merck, Darmstadt, Germany; final concentration, ~100 mM). Sample preparation of the engineered plant product isolate (putative 5-chloroindican) was carried out accordingly, while limited by the availability of only about 0.8 mg of the compound (final concentration, ~16 mM). The respective solutions were then transferred into 3 mm NMR tubes (Wilmad Labglass, Vineland, USA) for measurement.

NMR experiments were recorded on a Bruker Avance III HD 700 MHz spectrometer with a 5 mm quadruple channel inverse CryoProbe (1 H/19 F-31P/13 C/15 N/2 H) equipped with a z-gradient coil with maximum amplitude of 53 G/cm (Bruker BioSpin GmbH, Rheinstetten, Germany). Temperature was regulated with a BCU-II chiller and the sensor was corrected against a 99.8% methanol-*d*_4_ sample (Findeisen et al. 2007). All measurements were carried out at 300 K. The Bruker TopSpin 3.2pl6 software was used for data acquisition, while their processing was conducted in TopSpin 3.5pl6, DynamicsCenter 2.4.4 (Bruker BioSpin GmbH, Rheinstetten, Germany), and MestreNova 10.0.1 (MestreLab Research, Santiago de Compostela, Spain). Spectrum referencing was performed by setting the residual solvent peak of methanol to 3.310 ppm. All other spectra were referenced according to the Ξ scale. All samples were subjected to the same set of experiments. Unless otherwise noted, the default pulse sequences and parameters from the Bruker library were used: ^1^H, ^13^C, ^1^H-^1^H CLIP-COSY (sequence according to Koos et al., 2016; (Koos et al. 2016), 40 ms mixing), ^1^H-^1^H NOESY (zero-quantum suppression with a Thrippleton-Keeler filter (Thrippleton and Keeler 2003) 300 ms mixing), ^1^H-^13^C HSQC (transfer delay corresponding to 145 Hz), ^1^H-^13^C HMBC (long-range delay corresponding to 8 Hz), ^1^H-^15^N HMBC (long-range delay corresponding to 5 Hz), ^1^H DOSY (stimulated echo, bipolar gradient pairs, δ/2 = 1 ms, Δ = 100 ms, 32 gradient steps linearly spaced between 2% and 98%). Additionally, for the plant extract, ^1^H PSYCHE-iDOSY (sequence according to Foroozandeh et al. (2016)) (Foroozandeh et al. 2016) was recorded (PSYCHE: 20°Flip angle, 10 kHz bandwidth of the saltire chirp pulses with 15 ms duration, 20 increments with a spectral window of 70 Hz in the pure shift dimension; DOSY: δ = 1.5 ms, Δ = 100 ms, 32 gradient steps linearly spaced between 2% and 98%; pseudo 3D data reconstructed to pseudo 2D data with the *pshift* macro from http://nmr.chemistry.manchester.ac.uk and subsequently fitted with DynamicsCenter).

## Abbreviations

(i)DOSY: (integrated into) diffusion ordered spectroscopy
2A6: human cytochrome P450 monooxygenase 2A6
BFP: blue fluorescent protein
Bx1: indole synthase from *Zea mays*
CaMV: cauliflower mosaic virus
CLIP-COSY: clean in-phase correlation spectroscopy
cp: chloroplast
DW: dry weight
ER: endoplasmic reticulum
ESI: electrospray ionization
EV: empty vector
Fd: ferredoxin
GB2.0: GoldenBraid 2.0 modular cloning system
GT: glucosyltransferase
HCl: hydrochloric acid
HMBC: heteronuclear multiple bond correlation
HPLC: high performance liquid chromatography
HSQC: heteronuclear single quantum correlation
KBr: potassium bromide
*m/z*: mass-to-charge ratio
MIA(s): monoterpenoid indole alkaloid(s)
MS: mass spectrometry
NADH: nicotinamide adenine dinucleotide (reduced form)
NMR: nuclear magnetic resonance
NOE(SY): nuclear Overhauser enhancement (spectroscopy)
Nos: nopaline synthase
OD: optical density
P450: cytochrome P450 monooxygenase
PCR: polymerase chain reaction
PrnA: tryptophan 7-halogenase from *Pseudomonas fluorescens*
PSYCHE: pure shift yielded by chirp excitation
PyrH: tryptophan 5-halogenase from *Streptomyces rugosporus*
RebF: flavin reductase from *Lechevalieria aerocolonigenes*
RebH: tryptophan 7-halogenase from *L. aerocolonigenes*
*R_f_*: retention factor
RT: room temperature
STE: stimulated echo
SttH: tryptophan 6-halogenase from *Streptomyces toxitricini*
ThaL: tryptophan 6-halogenase from *Streptomyces albogriseolus*
TLC: thin layer chromatography
TnaA: tryptophanase from *Escherichia coli*
*t_R_*: retention time
Trp: tryptophan
TS: tryptophan synthase
TU: transcriptional unit
UPLC: ultra performance liquid chromatography
X-Gal: 5-bromo-4-chloro-indolyl-β-D-galactopyranoside
YFP: yellow fluorescent protein

